# *In vitro* effects of tensile strain on cancer-associated and matched normal fibroblasts derived from oral squamous cell carcinoma: an exploratory study

**DOI:** 10.64898/2026.07.11.737904

**Authors:** Nadine Wiesmann-Imilowski, Sebahat Kaya, Andressa V. B. Nogueira, Victoria Langer, Stefanie Zimmer, Johannes Mockenhaupt, Johannes U. Mayer, James Deschner, Jürgen Brieger, Peer W. Kämmerer

**Author notes:** **Corresponding author** Peer W. Kämmerer, Department of Oral and Maxillofacial Surgery, University Medical Center Mainz, Augustusplatz 2, 55131 Mainz, Germany.

## Abstract

**Background:** Mechanical forces, particularly tensile stress, influence tumor progression by modulating cancer-associated fibroblast (CAF) behaviour and extracellular matrix remodelling, yet their role in oral cancer remains insufficiently defined. To our knowledge, this is the first study investigating *in vitro* tensile stress responses in CAF and normal fibroblasts (NF) derived from oral squamous cell carcinoma (OSCC).

**Objective:** To assess how tensile stress affects fibroblast gap closure, proliferation, metabolic activity, and paracrine signalling relevant to tumor-stroma interactions.

**Methods:** CAFs and NFs were isolated from OSCC tissue and matched healthy mucosa and exposed to cyclic tensile strain (3%, 0.02 Hz, 96 h) using the FX-6000T™ system. Gap closure was assessed by wound-healing assay, proliferation by cell counting, metabolic activity by AlamarBlue™, and paracrine effects on A549 tumor cell gap closure using fibroblast-conditioned supernatants.

**Results:** Tensile loading significantly increased CAF gap closure capacity compared with both stimulated NFs (p<0.0001) and unstimulated CAF controls (p=0.0167). Metabolic activity showed a non-significant trend toward higher values in CAFs. Proliferation did not differ between groups up to 48 h, arguing against a major early contribution of proliferation to the observed group differences in the fibroblast scratch assays; however, because proliferation was not inhibited and was not quantified beyond 48 h, later time points should be interpreted conservatively as composite gap closure. Supernatants from mechanically stimulated CAFs increased A549 gap closure by ∼40% compared with non-stimulated controls (p=0.0485), whereas stimulated NFs did not enhance the tumor cell’s capacity to close the cell-free gap.

**Conclusions:** Tensile strain was associated with increased fibroblast gap closure and enhanced gap-closure–promoting paracrine effects of oral CAFs under the conditions tested. These exploratory findings support the concept that biomechanical cues can modulate CAF functional behaviour in OSCC. Given the limited cohort size and inter-patient variability, these results should be interpreted as hypothesis-generating and require confirmation in larger studies. Mechanistic pathways were not interrogated in this study and should be addressed in future work using more complex tumor-stroma models.

**Graphical abstract:** 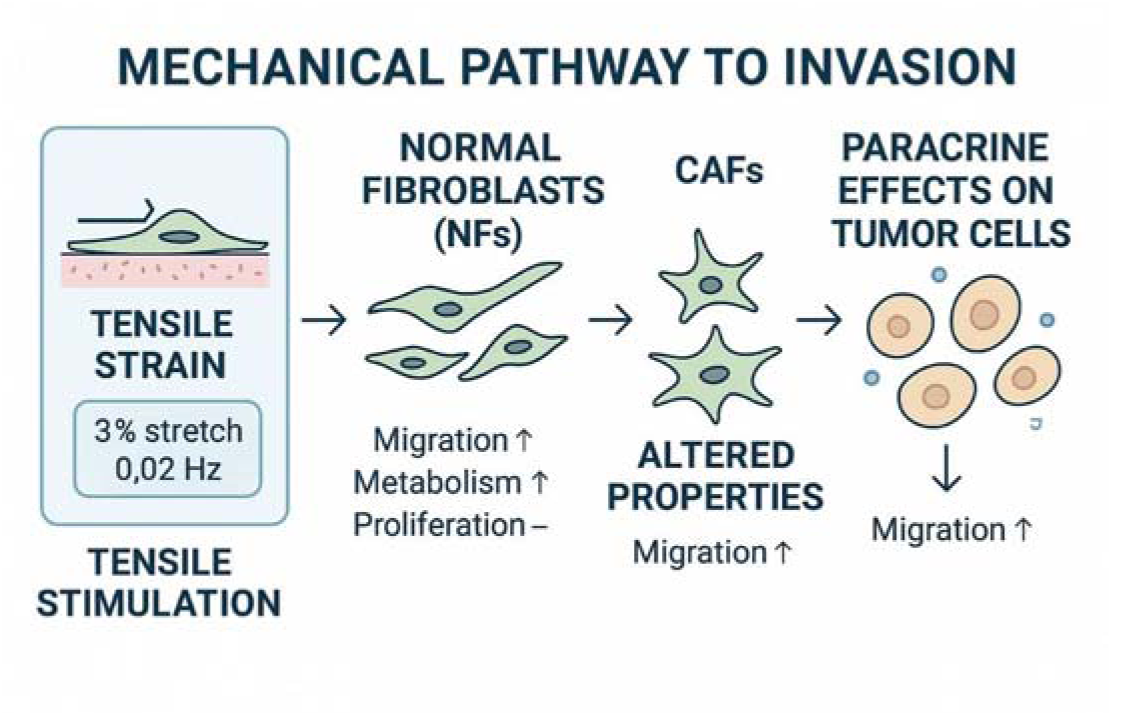

## Background

Tumors profoundly reshape their surrounding microenvironment, promoting growth and invasion through interactions between malignant cells, stromal components, and the extracellular matrix (ECM) [1–4]. Among these alterations, biomechanical changes within the tumor microenvironment (TME) have gained increasing attention, although their contribution to tumor behaviour remains incompletely understood [5].

Oral squamous cell carcinoma (OSCC) is uniquely exposed to mechanical loading due to continuous motion during speech, mastication, and swallowing, subjecting tumor tissues to compression, shear, and tensile forces [6, 7]. Matrix stiffness has been linked to mechanotransduction pathways involving factors such as YAP/TAZ, integrin signalling, and mechanosensitive ion channels, which may contribute to aggressive tumor phenotypes [8–11]. Mechanical stress and stiffness also influence oncogenic signalling in tongue carcinoma models [12].

Fibroblasts are central mediators of these mechanobiological interactions. They facilitate oral cancer invasion in collagen matrices, and increased matrix stiffness in OSCC correlates with recurrence and poor survival [13, 14]. Mechanotransduction pathways, including the cGAS-STING axis, have been implicated in fibroblast activation and ECM remodelling in several tumor contexts [15]. Cancer cells further the conversion of normal fibroblasts (NFs) into cancer-associated fibroblasts (CAFs) through mechanical cues and soluble mediators such as transforming growth factor-α and platelet-derived growth factors [16, 17]. CAFs enhance ECM deposition and lysyl oxidase (LOX)-mediated cross-linking, driving epithelial-mesenchymal transition (EMT) and promoting invasion. Mechanical cues have been linked to epithelial plasticity, including EMT-like programs, in several tumor settings [14, 15]. Their abundance increases with tumor stage in tongue carcinoma [18], and ECM stiffening further modulates migration, proliferation, and phenotypic plasticity in oral cancer cells [19–22]. Tumor stiffness also carries prognostic value in other cancers [23].

Despite these insights, the role of tensile forces, particularly relevant at invasive tumor fronts, remains poorly characterized. Mechanical stimulation has been implicated in oral carcinogenesis [24, 25], yet the effects of tensile strain on primary oral CAFs and matched NFs have not been systematically examined.

This pilot study investigated how cyclic tensile strain influences gap closure and paracrine signalling in CAFs and NFs derived from OSCC. Understanding these responses may help generate testable hypotheses on how mechanical cues shape CAF behaviour in OSCC and inform future mechanistic studies. The present work focuses on fibroblast functional responses and paracrine effects; mechanistic endpoints such as EMT were not assessed and are beyond the scope of this exploratory study.

## Materials and Methods

### Cell Isolation and Cell Culture

Primary fibroblasts were isolated from tumor tissue and matched healthy oral mucosa of patients with OSCC treated at the Department of Oral and Maxillofacial Surgery - Plastic Surgery, University Medical Center Mainz. Tumor samples and corresponding healthy mucosa were collected during surgical resection. Healthy biopsies were obtained from clinically normal oral mucosa at the site farthest from the resection margin, allowing direct comparison between cancer-associated fibroblasts (CAFs) and normal fibroblasts (NFs) from the same patient.

All procedures involving human tissue were performed in accordance with the Declaration of Helsinki and approved by the Ethics Committee of the State Medical Association of Rhineland-Palatinate (2022-16424_3). Written informed consent was obtained from all donors. All samples were pseudonymized prior to processing.

Biopsies were rinsed in phosphate-buffered saline (PBS; Merck, Darmstadt, Germany) supplemented with Amphotericin B (AMPHO-MORONAL; Dermapharm AG, Grünwald, Germany) to prevent fungal contamination. Tissue pieces were cut into fragments of approximately 2–3 mm³ and transferred into 6-well plates. To facilitate adherence, fragments were incubated without medium for 7 min at 37 °C. They were then covered with Dulbecco’s Modified Eagle Medium (DMEM; Gibco®, Life Technologies™, Paisley, UK) supplemented with 10% fetal bovine serum (FBS), 1% penicillin–streptomycin–neomycin, and 1% L-glutamine (Sigma-Aldrich, Steinheim, Germany). Cultures were maintained at 37 °C and 5% CO_2_ and monitored regularly. Medium was changed as needed until fibroblast outgrowth became apparent.

Once cultures reached 80–90% confluence, cells were detached using Accutase® (Thermo Fisher Scientific, Waltham, MA, USA) and transferred to culture flasks. CAFs and NFs were expanded up to passage 2. Mycoplasma testing was performed before the second passage. All experiments were conducted using cells in the same passage in which characterization was completed (passages 3–4).

For paracrine signalling experiments, A549 non-small cell lung carcinoma cells (DSMZ, Braunschweig, Germany) were used as a standardized and robust gap-closure readout model to quantify migration-related responses to fibroblast-conditioned media [26–28]. As A549 cells are not OSCC-derived, implications for OSCC-specific tumour–stroma interactions are addressed in the Discussion. Cells were cultured in DMEM/Ham’s F12 (Sigma-Aldrich, St. Louis, MO, USA) supplemented with 5% bovine calf serum (VWR Seradigm, Cat. No. 10158-358) and 1% penicillin–streptomycin (100 U/mL and 100 mg/mL, respectively) at 37 °C and 5% CO_₂_.

### Characterization of CAFs and NFs

Fibroblast outgrowth was first verified morphologically. The identity of cancer-associated fibroblasts (CAFs) and normal fibroblasts (NFs) was then confirmed by qPCR using a PrimePCR™ Custom 96-well Plate (Bio-Rad Laboratories Inc., Hercules, USA) containing predesigned and validated PrimePCR™ Assays (Table 1). Reference gene suitability was evaluated using the PrimePCR™ Pathway Plate “Reference Genes H96, human” (Bio-Rad Laboratories), and TFRC was selected.

**Table 1:**
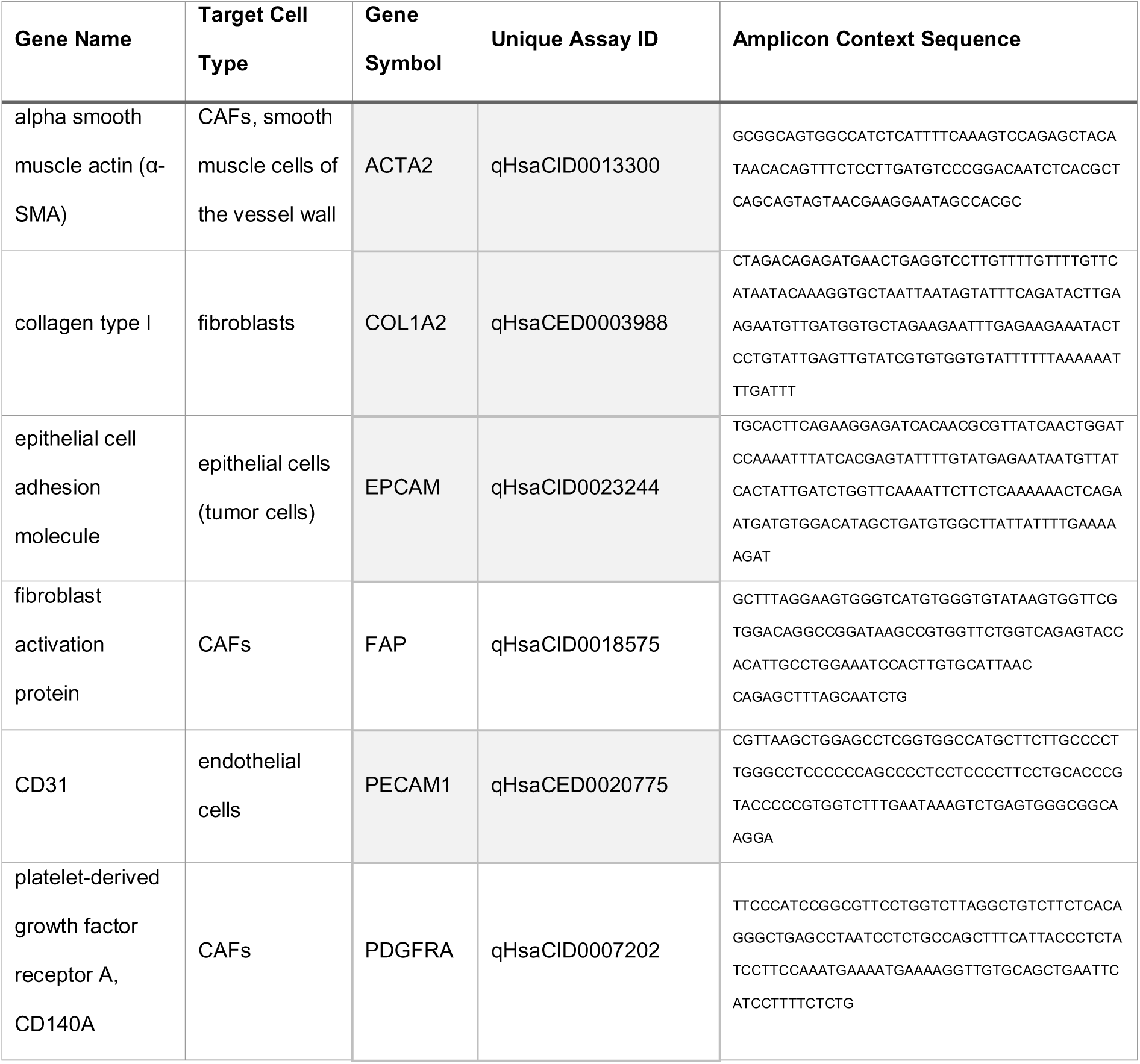
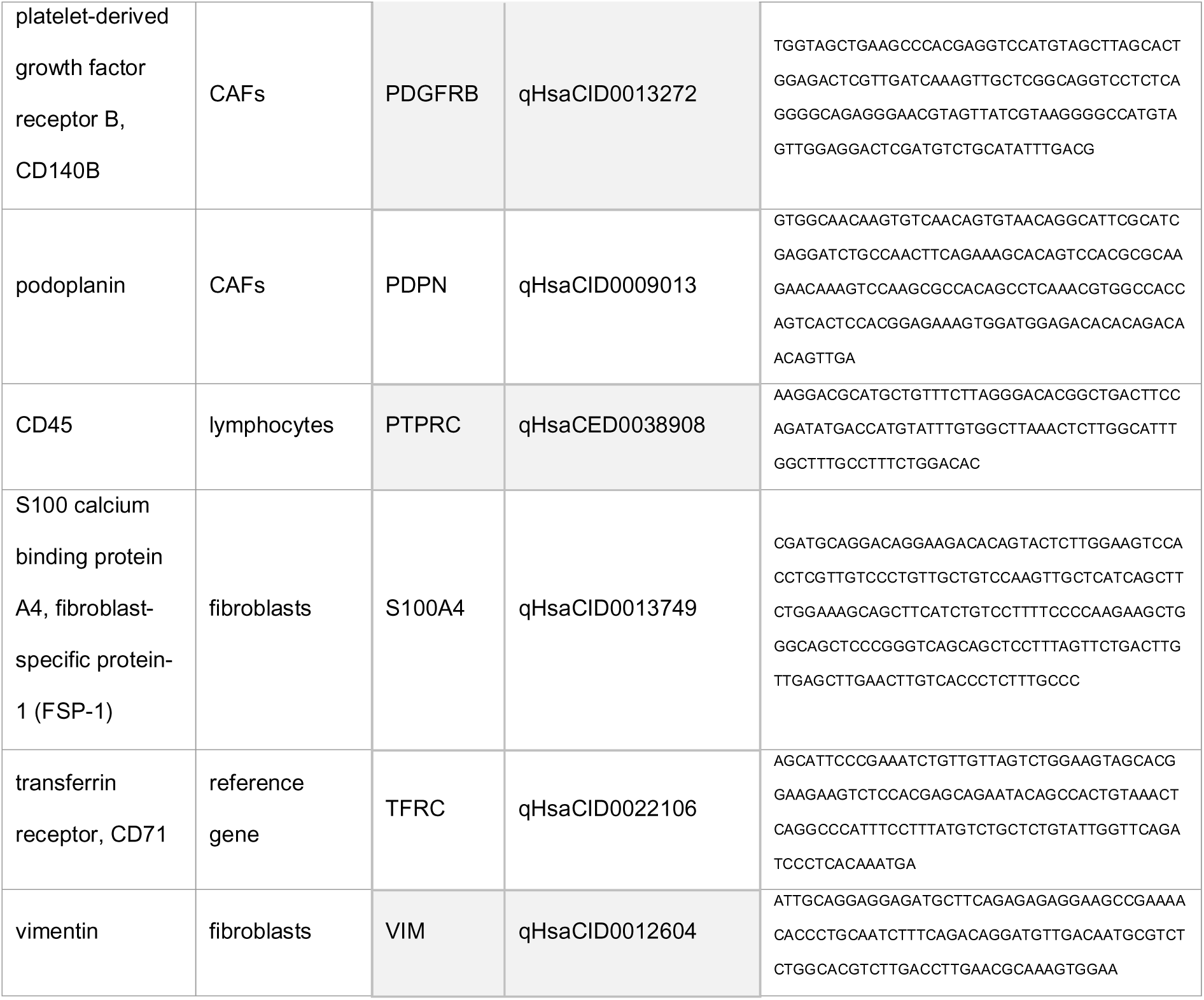
Marker genes used for identification and characterization of NFs and CAFs via qPCR.

To exclude contamination by other cell types, the absence of EPCAM (epithelial/tumor cells), CD31 (endothelial cells), and CD45 (lymphocytes) was confirmed. Fibroblast identity was assessed via expression of vimentin, collagen type I, and S100A4. CAF status was determined by expression of established CAF markers including α-SMA, PDPN, PDGFRA, PDGFRB, and FAP. Because fibroblasts and CAFs are heterogeneous, a multi-marker panel was applied to capture a broad spectrum of CAF phenotypes [30–32]. Five patient-matched CAF/NF pairs were included in this pilot study.

As a representative orthogonal protein-level validation of the transcription-based classification, α-SMA expression was evaluated in one CAF/NF pair by immunocytochemistry. Given the known heterogeneity of CAF phenotypes, this representative α-SMA staining was not used as the sole basis for CAF classification, which relied primarily on the multi-marker qPCR panel together with exclusion of EPCAM-, CD31-, and CD45-positive contamination. For this purpose, 10,000 cells were seeded onto sterile glass coverslips (VWR International GmbH, Darmstadt, Germany) placed in 6-well plates.

After four days, cells were washed twice with PBS (Merck, Darmstadt, Germany) and fixed in 4% paraformaldehyde (PFA; Thermo Fisher Scientific, Waltham, MA, USA) for 20 min. Cells were washed with TBST20 (Tris-buffered saline, 0.05% Tween-20; Sigma-Aldrich Chemie GmbH, Steinheim, Germany) and endogenous peroxidase activity was blocked using methanol/hydrogen peroxide (0.6% H_₂_O_₂_ in MeOH; Merck) for 18 min. Cells were rinsed in distilled water and washed twice with TBST20 before blocking for 45 min with 10% normal goat serum (Vector Laboratories, Burlingame, CA, USA) in 1% BSA/PBS (Sigma-Aldrich).

Primary antibody incubation was performed overnight at 4°C using anti-α-SMA (mouse monoclonal, clone 1A4, #A2547; 1:1000 in 1% BSA/PBS; Sigma-Aldrich Chemie GmbH). The next day, cells were washed with TBST20 containing 400 mM NaCl (Sigma-Aldrich) followed by TBST20 alone. Secondary antibody (goat anti-mouse HRP, #ab6789; 1:250 in PBS; Abcam, Cambridge, UK) was applied for 1 h at room temperature. After two further TBST20 washes, immunoreactivity was visualized using DAB substrate (Zytomed Systems, Berlin, Germany; 10 min). Nuclei were counterstained with hematoxylin (5 min; Merck), rinsed in tap water, and coverslips were mounted using Aqueous Mount (Zytomed Systems). Negative controls lacking primary antibody were processed in parallel.

### Immunohistochemical Analysis of Patient Tissues for CAF Isolation

Formalin-fixed, paraffin-embedded (FFPE) sections were deparaffinized in xylene and rehydrated through graded ethanol (100%, 96%, 70%, 50%; 3 min each) before immersion in distilled water (3 min). Heat-induced epitope retrieval was performed in Dako Target Retrieval Solution, Tris/EDTA, pH 9 (Dako/Agilent, Santa Clara, USA) for 20 min in a steamer, followed by cooling under running water and one wash in EnVision™ FLEX Wash Buffer (5 min).

Immunostaining was performed on an Epredia™ Lab Vision™ Autostainer 480S (Epredia, Portsmouth, NH, USA) using the EnVision™ FLEX HRP/DAB detection system (kit K8010, Dako/Agilent) according to the manufacturer’s instructions. The primary antibody mouse anti-α-smooth muscle actin, clone 1A4, ready-to-use (REF IR611, Dako/Agilent), was applied without dilution.

Chromogenic visualization was achieved using DAB from the EnVision™ FLEX system. Slides were counterstained with Dako Hematoxylin (REF 2020), blued in tap water, and washed in distilled water (1 × 5 min). Sections were dehydrated through graded ethanol, cleared in xylene, and coverslipped using Entellan® mounting medium (Merck, Darmstadt, Germany).

### Mechanical Loading of the Fibroblasts

To simulate mechanical stress *in vitro*, CAFs and NFs (n = 5 each) were subjected to cyclic tensile strain for up to 4 days using the FX-6000T™ Tension System (Flexcell® International Corporation, Burlington, NC, USA), as previously described [29–31]. For this purpose, 200,000 fibroblasts per wellwere seeded onto 6-well BioFlex® Culture Plates with collagen-coated silicone membranes (Flexcell® International Corporation). Cells were allowed to adhere overnight.

The BioFlex® plates were then mounted in the FX-6000T™ device (Figure 1), and fibroblasts were exposed to 3% cyclic tensile strain at a frequency of 0.02 Hz for 96 h. This frequency corresponds to the approximate physiological swallowing rate of once per minute [32]. The strain amplitude (3%) and stimulation duration (96 h) were selected as a low-to-moderate cyclic tensile regimen based on literature precedent and on preliminary feasibility optimization for sustained loading of primary oral fibroblasts. In fibroblast mechanobiology, both magnitude and duration are known to influence the cellular response [33], and oral/periodontal loading studies have used low strain magnitudes including 3% as well as multi-day exposure windows [34]. Direct physiological quantification of tensile strain magnitudes in OSCC-associated oral soft tissues is currently limited; therefore, parameter selection was guided by precedent and feasibility rather than direct in vivo measurements. In our preliminary experiments, different strain magnitudes and stimulation durations were explored, and the final regimen of 3% strain, 0.02 Hz, and 96 h was selected because it enabled reproducible multi-day stimulation under stable culture conditions and was considered suitable for assessing chronic functional responses in patient-derived fibroblasts. Control fibroblasts were seeded onto identical BioFlex® plates and maintained in the same incubator under identical environmental conditions, but without mechanical loading.

**Figure 1:**
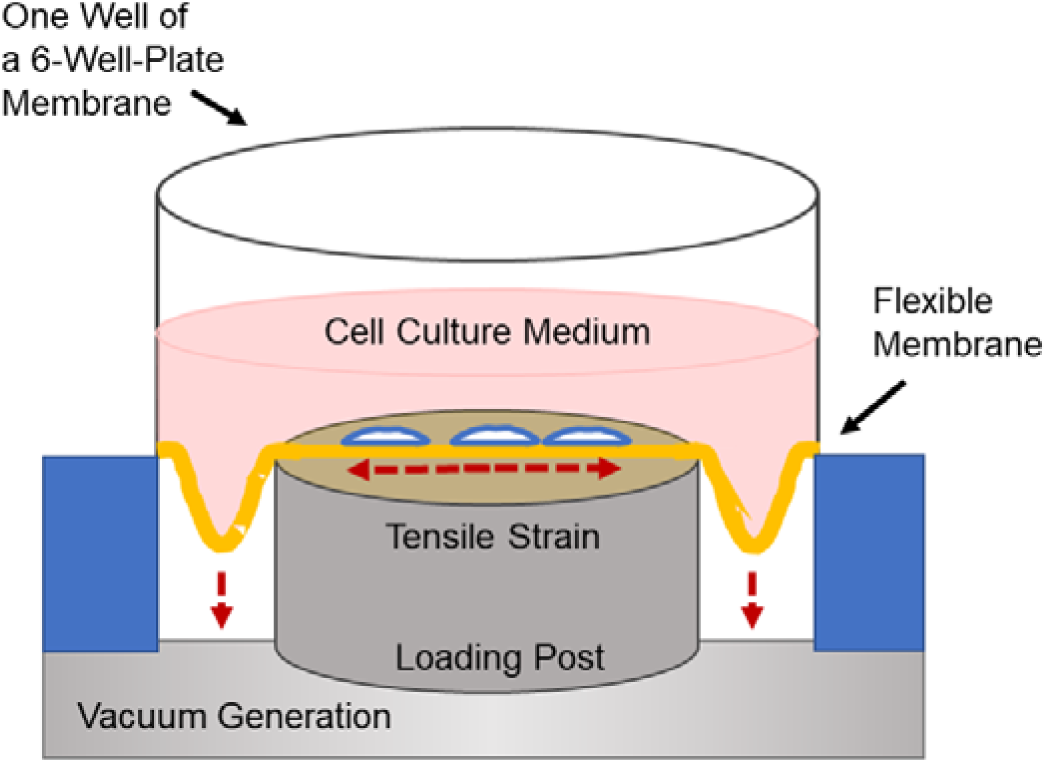
Schematic representation of the device for application of tensile strain on adherent cells *in vitro*. Fibroblasts were seeded on the BioFlex^®^ cell culture plate and were given time to adhere overnight. The plate was then placed in a stretching device that applied a controlled tensile strain on the cells. This system includes a flexible silicone membrane at the base of the cell culture plate, which is stretched by the downward movement by vacuum generation. This movement generates a controlled tensile strain on the cells, simulating similar conditions *in-vivo*.

After completion of mechanical stimulation, cell metabolic activity of the cells was assessed. Supernatants were collected and stored for subsequent medium-transfer experiments to investigate paracrine effects on tumor cell’s capacity of gap closure. Fibroblasts were then harvested and reseeded for evaluation of migratory and proliferative properties.

### Analysis of Cellular Metabolic Activity

Cellular metabolic activity after mechanical loading was assessed using the AlamarBlue™ Cell Viability Reagent (ThermoFisher Scientific, Waltham, MA, USA) according to the manufacturer’s instructions and as previously described [35, 36]. Following mechanical stimulation, the culture medium in the BioFlex® plates was replaced with fresh medium containing 10% AlamarBlue™ reagent. Fibroblasts were incubated for 4 h at 37 °C, after which relative fluorescence was measured using a Fluoroskan Ascent Microplate Reader (ThermoFisher Scientific, Waltham, MA, USA) with 538 nm excitation and 600 nm emission filters.

### Wound Healing Assay

Fibroblast migration in form of gap closure was assessed using a wound healing assay as previously described [35, 37] with culture inserts in 35 mm dishes (ibidi®, Munich, Germany). After mechanical loading, fibroblasts were harvested, manually counted, and seeded into each well of the culture insert at 70 µl of a suspension containing 400,000 cells. Following overnight adhesion, inserts were removed to create a standardized cell-free gap. Because proliferation was not pharmacologically inhibited (e.g., with Mitomycin C), the readout generally reflects gap closure as a composite of cell migration and proliferation [38]. To obtain an indication of whether gap closure was due to proliferation or migration, proliferation was always assessed in parallel.

Gap closure was documented by light microscopy at 0 , 24 , 48 , 72 , and 96 h. Quantitative analysis was performed using T-scratch software (https://www.cselab.ethz.ch/software). Gap closure was expressed as the relative open area, defined as 100% at 0 h and decreasing as the gap closed over time.

For comparative analysis, untreated NFs served as the reference group (100% at all time points). CAFs and NFs were always compared as matched pairs from the same patient. A total of n = 5 CAF/NF pairs was analyzed.

### Assessment of Cell Proliferation

Cell proliferation was quantified using harvested fibroblasts. The cell suspension was diluted 1:15 in culture medium, and 1 ml of the diluted suspension was seeded outside the culture insert. Proliferation was documented by light microscopy at 0 h, 24 h, and 48 h.

For analysis, fibroblasts were manually counted in four representative fields per time point using the ImageJ Fiji Cell Counter. The number of cells at 0 h was set to 100%, and subsequent values were expressed relative to this baseline.

### Analysis of the Paracrine Effects of Fibroblasts on Tumor Cell Gap Closure

To minimize serum-derived effects, fibroblasts were cultured in medium containing 2% FCS and 1% antibiotics before mechanical loading. After completion of loading, supernatants from CAFs and NFs (mechanically loaded and untreated controls) were collected and applied to tumor cells.

Paracrine effects on tumor cell gap closure were assessed using a wound-healing assay as previously described [35, 37] with culture inserts in 35-mm dishes (ibidi®, Munich, Germany). Each well of the culture insert was filled with 70 µl of a suspension containing 500,000 A549 tumor cells, which were allowed to adhere overnight. After insert removal, a standardized cell-free gap was generated, and tumor cells were incubated with the respective fibroblast-conditioned supernatants. This assay was used as a functional readout of gap-closure-related paracrine activity and does not represent an OSCC-specific epithelial compartment.

Gap closure was documented by light microscopy at 0 , 24 , 48 , 72 , and 96 h. Quantitative analysis was performed using T-scratch software (https://www.cselab.ethz.ch/software). Gap closure was expressed as relative open area, defined as 100% at 0 h and decreasing as cells filled the gap.

Supernatants from **untreated NFs** served as the reference condition (100% at all time points). Conditioned media from matched CAFs and NFs, with or without mechanical loading, were compared across **four groups**:

1. NF control,
2. mechanically loaded NF,
3. CAF control,
4. mechanically loaded CAF.

To obtain an indication of whether gap closure was due to proliferation or migration, proliferation was always assessed in parallel.

### Statistical analysis

Data were analysed using GraphPad Prism (Version 6.01, GraphPad Software, La Jolla, CA, USA). Descriptive statistics are reported as mean ± standard deviation (SD). All experiments were performed using patient-matched CAF/NF pairs (n = 5 donors); where fewer pairs were available for a given assay, the corresponding n is indicated in the respective figure legend. Comparative analyses were performed to assess the effects of mechanical loading on fibroblast metabolic activity, migration/gap closure, and proliferation, as well as the paracrine influence of fibroblast-conditioned supernatants on tumor cell migration/gap closure. For metabolic activity, normally distributed paired data were evaluated using a two-tailed paired t-test, whereas non-parametric paired data were analysed with the Wilcoxon matched-pairs signed-rank test. Differences in time-course outcomes (gap closure and proliferation) were examined using two-way repeated-measures ANOVA (matching by patient) with Bonferroni post hoc corrections for multiple comparisons. A p-value < 0.05 was considered statistically significant (* p < 0.05, ** p < 0.01, *** p < 0.001, **** p < 0.0001).

## Results

Primary fibroblasts derived from OSCC tissue and matched healthy mucosa were successfully characterized as NFs and CAFs, respectively. Subsequently, their metabolic activity, migratory capacity, and proliferative behaviour were examined to evaluate the effects of cyclic tensile strain. In addition, supernatants from mechanically loaded and untreated fibroblasts were collected for medium-transfer experiments to assess the paracrine influence of CAFs and NFs on tumor cell gap closure.

### Characterization of CAFs and NFs

Paired CAF and NF samples from five OSCC patients (Table 2) were included in the analysis.

**Table 2:** Patient’s and tumor characteristics. This table summarises age, sex, tumor localization, and TNM classification of the patients included in this study. TNM staging follows the UICC/AJCC 8th edition. Abbreviations: pT, pathological tumor size; pN, pathological lymph node status (positive nodes/total nodes examined); L0, no lymphatic invasion; V0, no vascular invasion; Pn0, no perineural invasion; R0, resection with clear margins; G2, moderately differentiated tumor.

| Cell-ID | Age | Sex | Diagnose | TNM Classification |
| --- | --- | --- | --- | --- |
| CAF_P1 | 65 | male | OSCC, left lateral border of the tongue | pT3, pN1 (1/45), L0, V0, Pn0, G2, R0 |
| CAF_P2 | 81 | female | OSCC, left lateral border of the tongue | pT2, pN0 (0/33), L0, V0, Pn0, R0 |
| CAF_P3 | 60 | male | OSCC, left lateral border of the tongue | pT2, pN0(0/44), L0, V0, Pn0, G2, R0 |
| CAF_P4 | 58 | male | OSCC, right mandibular angle | pT4a, G2, pN0 (0/44), L0, V0, Pn0, R0 |
| CAF_P5 | 60 | female | OSCC, right lateral border of the tongue | pT2, pN0 (0/20), L0, V0, Pn0, R0 |

To verify cell identity, mRNA expression of lineage-specific markers was assessed by qPCR. The absence of EPCAM, CD31, and CD45 confirmed that neither epithelial nor endothelial nor hematopoietic cells were present in the isolated cultures.

All samples expressed the examined fibroblast markers (Fig. 2), although expression levels varied substantially between patients, consistent with known inter-individual heterogeneity. Displaying NFs and CAFs side-by-side for each donor highlighted both inter-patient variability and the transcriptional differences between fibroblasts from healthy mucosa and tumor-derived fibroblasts.

**Figure 2:**
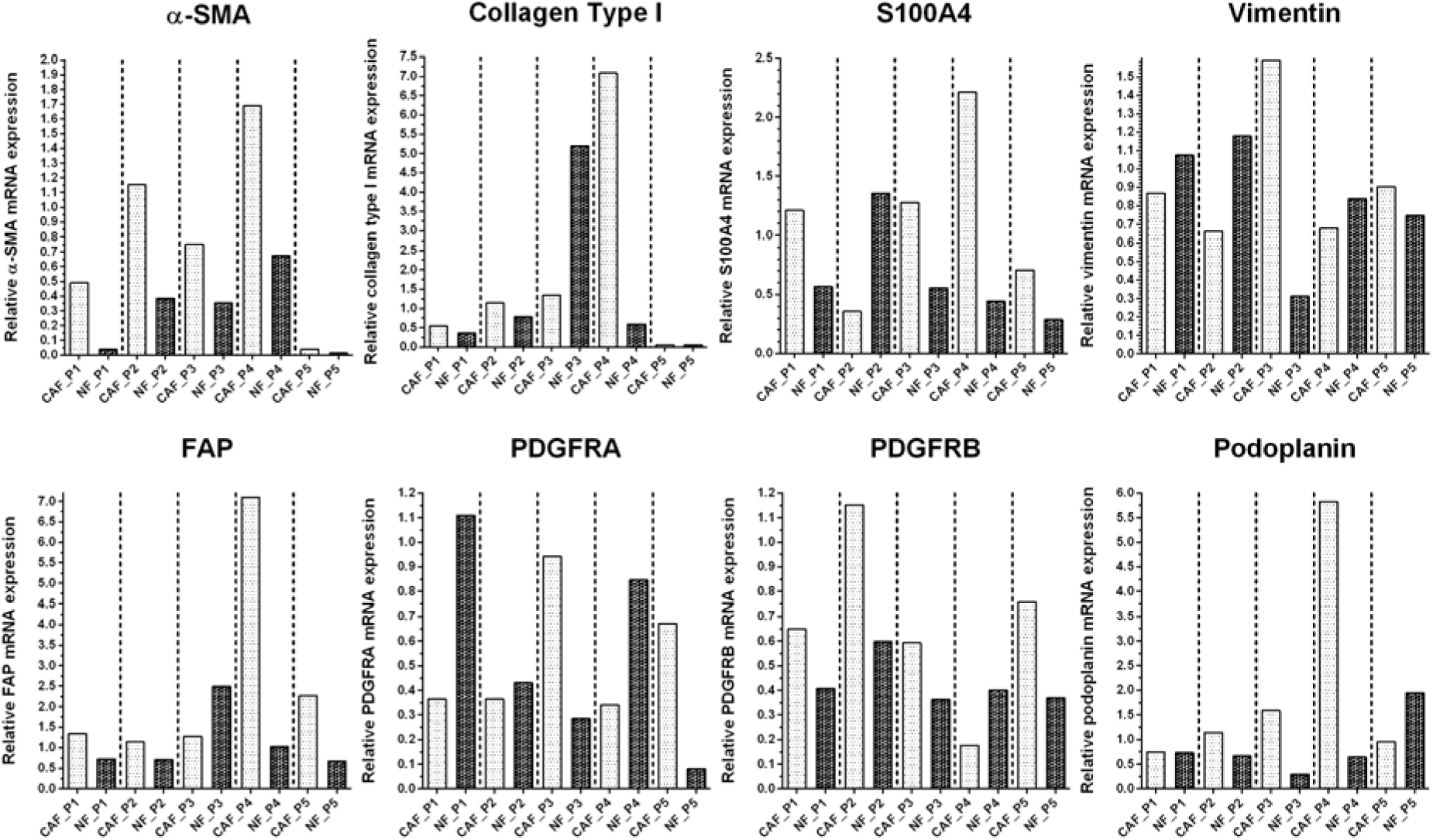
Characterization of CAF and NFs samples from five individual patients via mRNA expression of fibroblast-associated marker genes. Quantitative PCR analysis was performed for α-SMA, collagen type I, S100A4, vimentin, FAP, PDGFRA, PDGFRB, and podoplanin. CAFs and matched NFs were isolated from five patients (P1–P5). Bars represent relative mRNA expression levels of each marker in CAFs and NFs from the same patient.

To relate *in vitro* characteristics to *in-vivo* stromal features, α-SMA expression was examined in corresponding patient tissue specimens (Fig. 3). Tumor tissues consistently showed α-SMA positivity, but staining patterns differed between cases. In some samples, staining was confined to vascular smooth muscle cells, whereas others demonstrated α-SMA–positive CAFs within the tumor stroma with variable intensity. Examination of adjacent non-malignant tissue showed α-SMA expression predominantly restricted to vascular structures, even in cases with pronounced stromal α-SMA within the tumor core (e.g., P2, P4).

**Figure 3:**
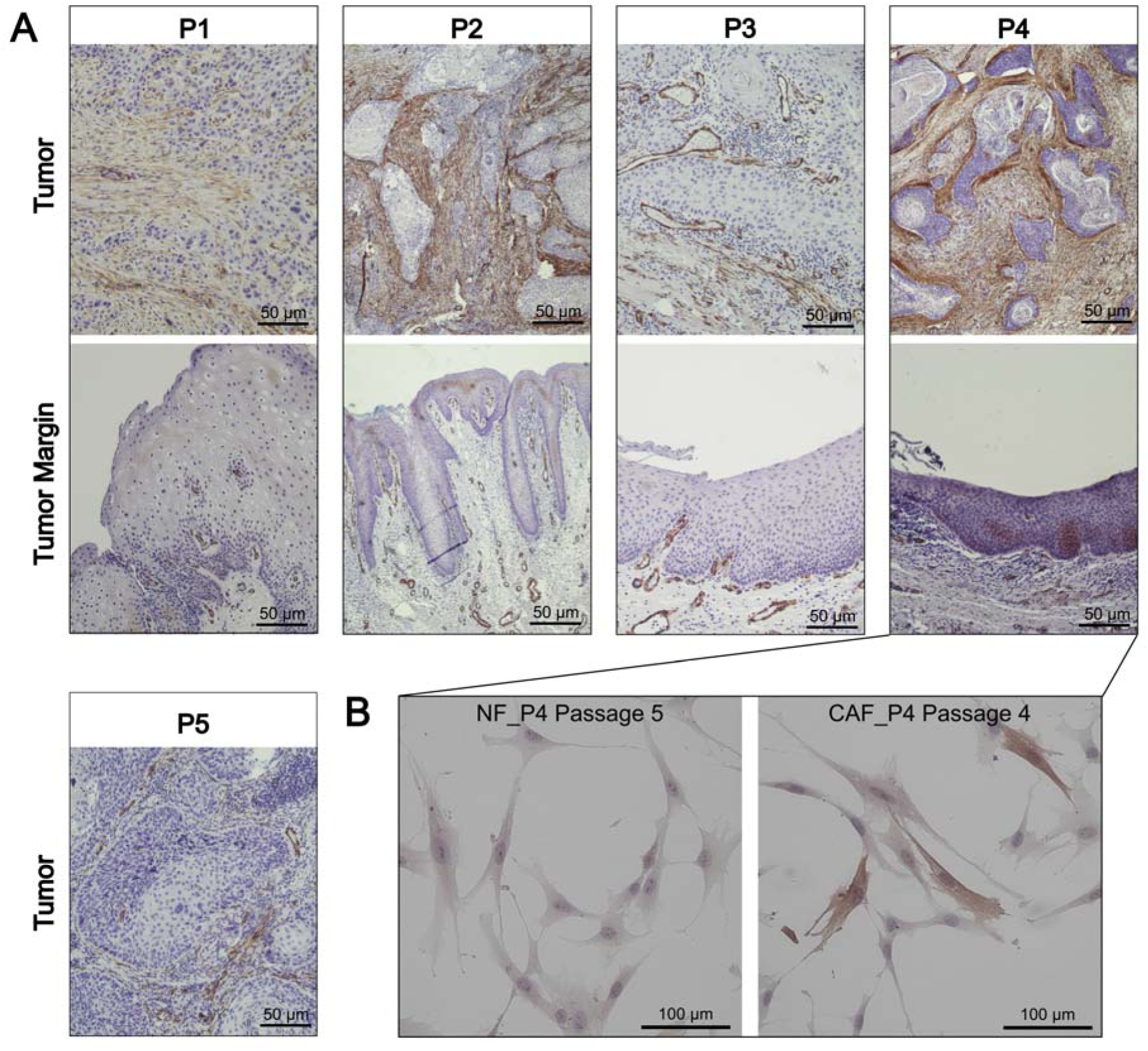
Immunohistochemical α-SMA Analysis of Patient Tissues for CAF Isolation and expression *in vitro*. (A) Representative area of the α-SMA immunohistochemical staining in tumor tissue and, where available, the corresponding tumor margins from patients P1–P5, whose tissue was used for fibroblast isolation. Variable α-SMA expression was observed in both tumor stroma, presumably in CAFs, and vascular smooth muscle cells, with inter-patient heterogeneity. In the tumor margins, α-SMA expression was predominantly restricted to vascular smooth muscle cells. (B) Immunocytochemical staining of fibroblasts isolated from patient P4, comparing NF (passage 5) and CAF (passage 4). CAFs showed stronger α-SMA expression compared to NFs, consistent with *in situ* findings.

The heterogeneity observed in tissue staining was reflected in the *in vitro* mRNA expression profiles of the isolated fibroblasts, supporting the notion that the cultured cells preserved key patient-specific features. Together with strict quality controls - including early-passage use and avoidance of freeze-thaw cycles - we considered the integrity of NF and CAF characteristics to be maintained for subsequent experiments.

### *In vitro* Wound Healing Assay

Fibroblasts from all groups exhibited a time-dependent reduction of the open scratch area, yet distinct differences were observed between NFs and CAFs and their mechanically stimulated counterparts (Fig. 4). Across all measured time points, CAFs showed significantly higher gap closure than matched NFs (24 h: p = 0.0256; 48–96 h: p < 0.0001). Tensile strain further enhanced gap closure in both cell types. After 96 h, mechanically stimulated NFs closed the gap significantly faster than unstimulated controls (p = 0.0089), and tensile stress similarly increased gap closure in CAFs compared with their untreated counterparts (p = 0.0167). Mechanically loaded CAFs displayed the fastest gap closure across all time points, suggesting a higher sensitivity of CAFs to tensile cues under the conditions tested.

**Figure 4:**
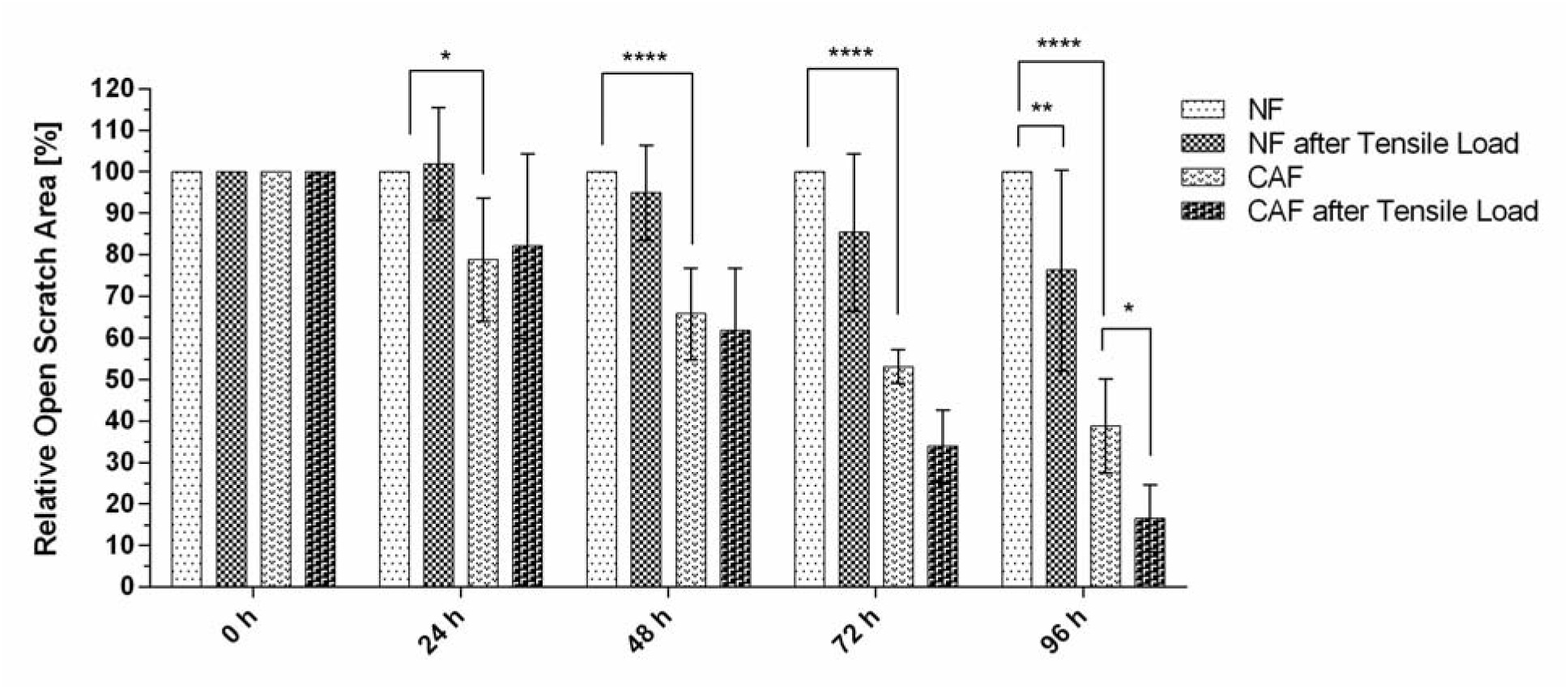
Gap closure of NFs and CAFs following mechanical loading. The migratory activity of the fibroblasts was represented as the relatively open area. An open area of 100% was assumed at the start time, which decreased over time in all investigated groups. CAFs exhibited an increased gap closure in comparison to NFs from the same patient. Tensile loading of the fibroblasts increased gap closure in both NFs and CAFs. Shown are mean values ± SD, the relative open area concerning the starting time, and the untreated NFs, which served as a control group. A two-way repeated-measures ANOVA (matching by patient) was performed to compare NFs and CAFs and their untreated and loaded counterparts, with Bonferroni correction for multiple comparisons (n = 5 patient-matched pairs).

As expected for primary human fibroblasts, considerable inter-patient variability in absolute gap closure rates was observed. Nevertheless, the relative differences between the experimental groups were largely consistent across all samples. Representative images from one patient (Fig. 5) illustrate the higher migratory activity of CAFs compared with NFs, as well as the additional increase induced by tensile loading.

**Figure 5:**
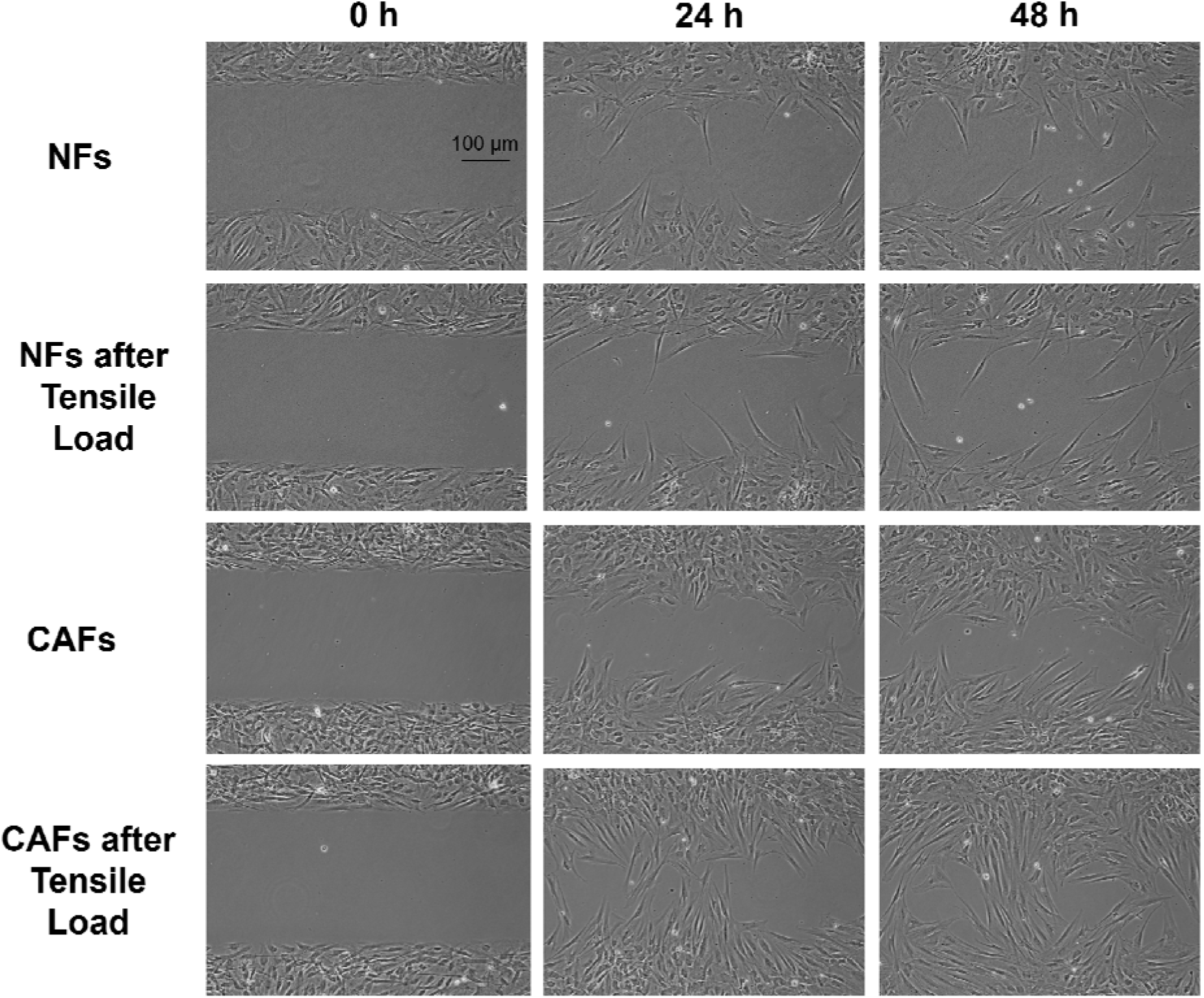
Representative depiction of the gap closure of NFs and CAFs of one set of patient samples following exposure to tensile load.

To further inform the interpretation of the gap-closure assay, proliferation was subsequently assessed in an independent assay.

### Proliferation Assay

No significant differences in fibroblast proliferation were detected between the groups at 0 h, 24 h, or 48 h (Fig. 6). At 48 h, unstimulated CAFs showed a non-significant increase in proliferation compared with unstimulated NFs (p = 0.4055). Conversely, mechanically stimulated CAFs displayed a modest reduction in proliferation relative to their unstimulated counterparts (p = 0.3496). However, substantial inter-patient variability among the primary fibroblast samples limited the ability to draw more definitive conclusions regarding proliferation behavior. Taken together, these data do not support a major differential proliferation effect up to 48 h. However, because proliferation was not inhibited in the scratch assay and was not quantified beyond 48 h, a contribution of proliferation to the 72–96 h wound-closure data cannot be excluded; these later time points should therefore be interpreted conservatively as composite gap closure.

**Figure 6:**
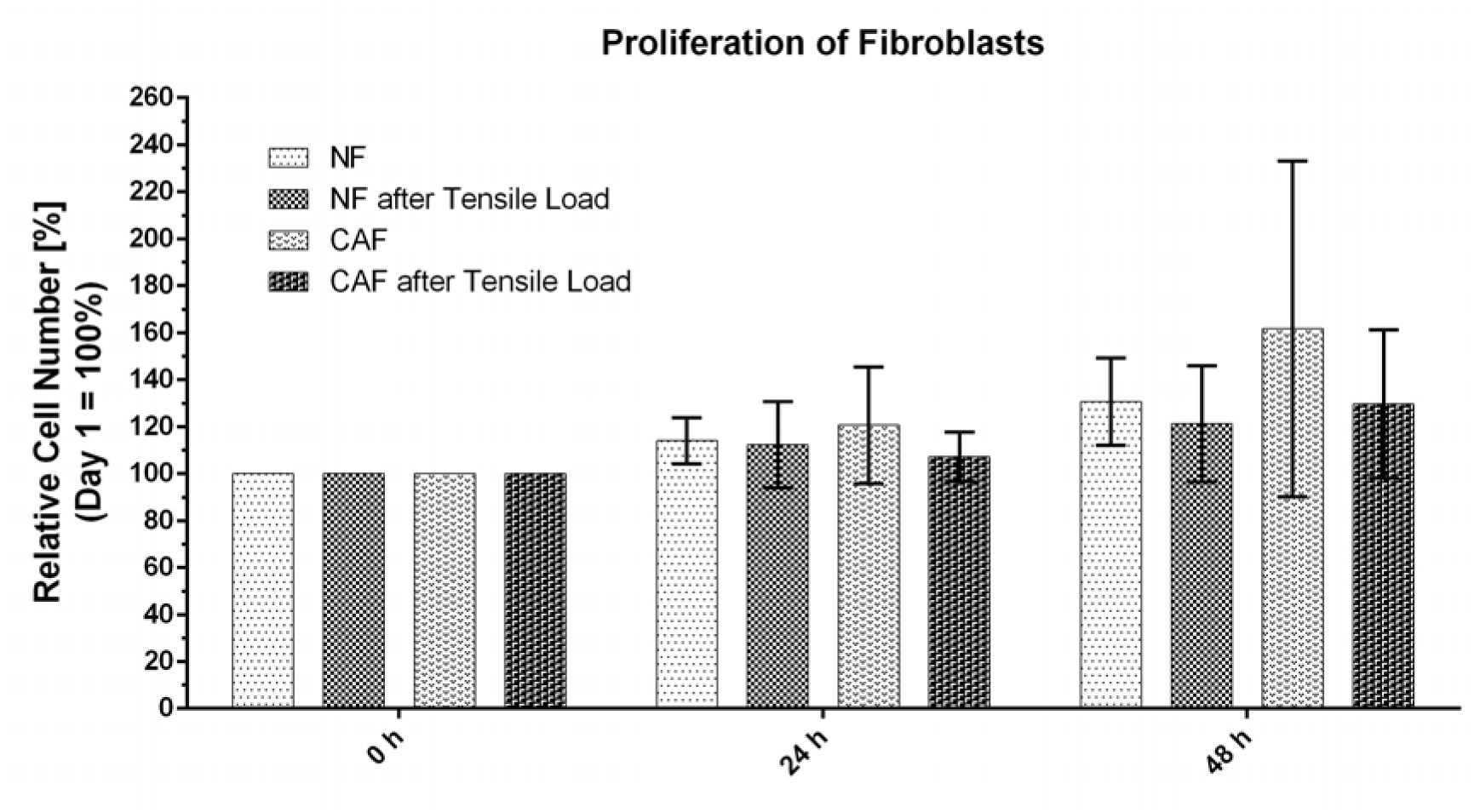
Proliferation of NFs and CAFs following mechanical loading. Significant differences in proliferation were not observed among the four fibroblast groups under investigation at 0 h, 24 h, and 48 h. Shown are mean values ± SD of the relative cell number as a measure for cell proliferation in relation to the starting time (Day 1 = 0 h = 100%). A two-way repeated-measures ANOVA (matching by patient) was performed comparing between NFs and CAFs and between the untreated and the loaded counterparts, with Bonferroni correction for multiple comparisons (n = 5 patient-matched pairs).

### Metabolic Activity

Metabolic activity was assessed immediately after tensile loading using a fluorescence-based assay. No statistically significant differences were observed between the experimental groups (Fig. 7). CAFs showed a non-significant trend toward higher metabolic activity compared with NFs (p = 0.1156). A slight non-significant decrease in metabolic activity was noted in mechanically stimulated CAFs relative to their unstimulated counterparts (p = 0.6791).

**Figure 7:**
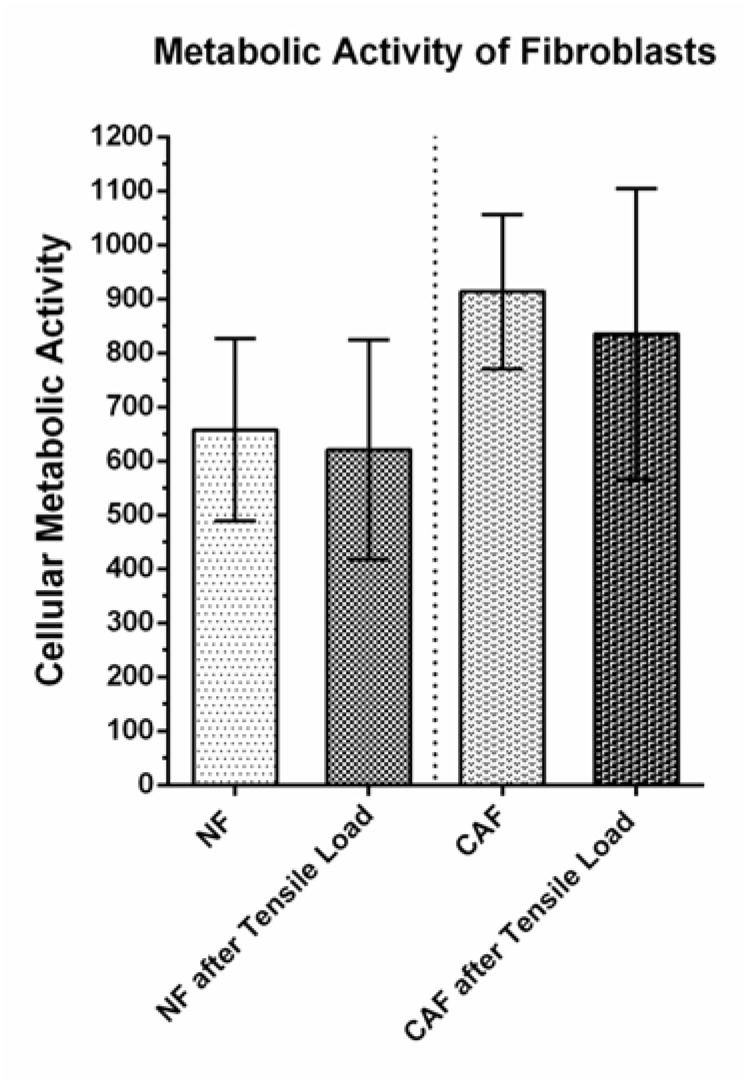
Cellular Metabolic Activity of NFs and CAFs following Mechanical Loading. No significant differences in the metabolic activity of CAFs and NFs were observed following tensile loading. However, there was a trend towards increased metabolic activity in CAFs compared to NFs. In contrast, a slightly reduced metabolic activity was observed in the group of stimulated CAFs compared to CAFs not subjected to tensile loading. Shown are mean values ± SD (patient-matched pairs; n as indicated in the dataset). A two-tailed paired t-test was performed comparing between NFs and CAFs and between the untreated and the loaded counterparts.

### Paracrine Effects of Fibroblasts on Tumor Cells

Paracrine influences of fibroblasts on tumor cell behaviour were assessed using A549 cells as a standardized gap-closure readout and incubating them with fresh supernatants from matched NFs and CAFs. The conditioned media had a notable impact on tumor cell gap closure (Fig. 8). After 24 h, a non-significant trend toward enhanced gap closure was observed in A549 cells treated with CAF supernatants compared with those treated with NF supernatants (p = 0.2312). This difference became significant at later time points, with CAF-derived supernatants promoting faster gap closure than NF-derived supernatants (48 h: p = 0.002; 72 h: p = 0.0134; 96 h: p < 0.0001).

**Figure 8:**
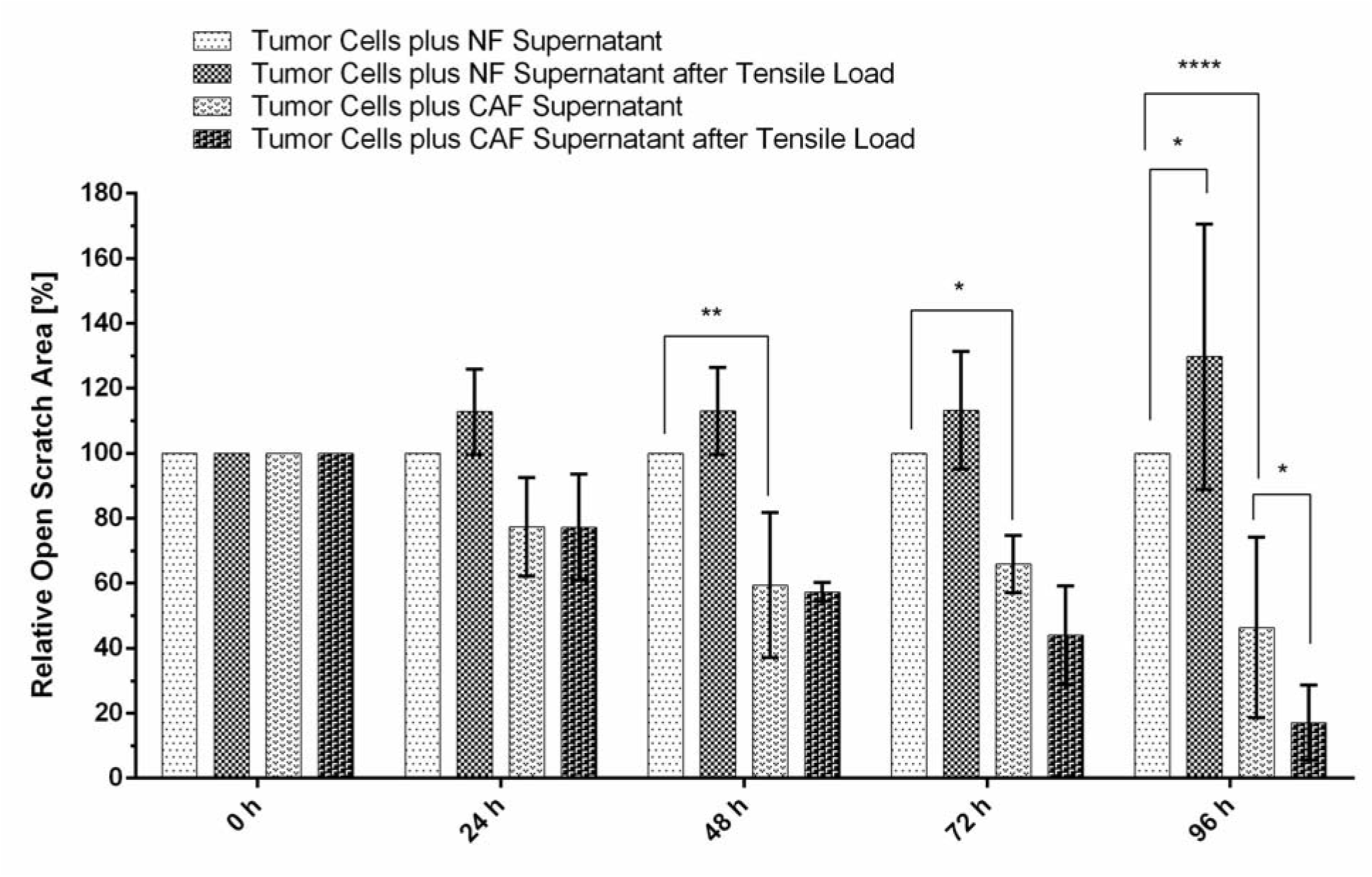
Paracrine Effects of Fibroblasts after Subjection to Mechanical Load on Tumor Cell Gap Closure. A549 cells were used as a standardized gap-closure readout and incubated with conditioned media from matched NFs and CAFs (unstimulated controls or after tensile loading). Gap closure is displayed as relative open area (0 h = 100%), normalized to A549 cells incubated with supernatant from unstimulated NFs (control). Conditioned media from CAFs increased tumor cell gap closure compared with NF supernatants. Supernatants from mechanically loaded CAFs further increased gap closure, whereas supernatants from mechanically loaded NFs reduced gap-closure–promoting activity compared with unstimulated NF controls. Shown are mean values ± SD. A two-way repeated-measures ANOVA (matching by patient) was performed comparing between NFs and CAFs and between the untreated and the loaded counterparts, with Bonferroni correction for multiple comparisons, n = 4 patient-matched pairs (* p < 0.05, ** p < 0.01, **** p < 0.0001).

Mechanical loading further modulated these paracrine effects. Supernatants from mechanically stimulated CAFs increased tumor cell gap closure at 96 h compared with unstimulated CAFs (p = 0.0485). Conversely, tensile loading reduced the gap-closure–promoting capacity of NF supernatants; conditioned media from loaded NFs caused significantly less tumor cell gap closure than media from unstimulated NFs at 96 h (p = 0.0435).

Inter-patient variability contributed to differences in response magnitude and relatively high standard deviations, consistent with the heterogeneity of primary fibroblast cultures. Nevertheless, all patient-derived pairs showed the same directional relationship: CAF supernatants consistently induced stronger tumor cell gap closure than NF supernatants, and mechanical loading amplified this effect specifically in CAFs. In addition to increased gap closure, A549 cells cultured with CAF-derived supernatants displayed altered migratory behavior: instead of advancing as a cohesive sheet, individual cells detached from the epithelial front and migrated as single cells. Representative images illustrating these effects for one patient sample are shown in Fig. 9.

**Figure 9:**
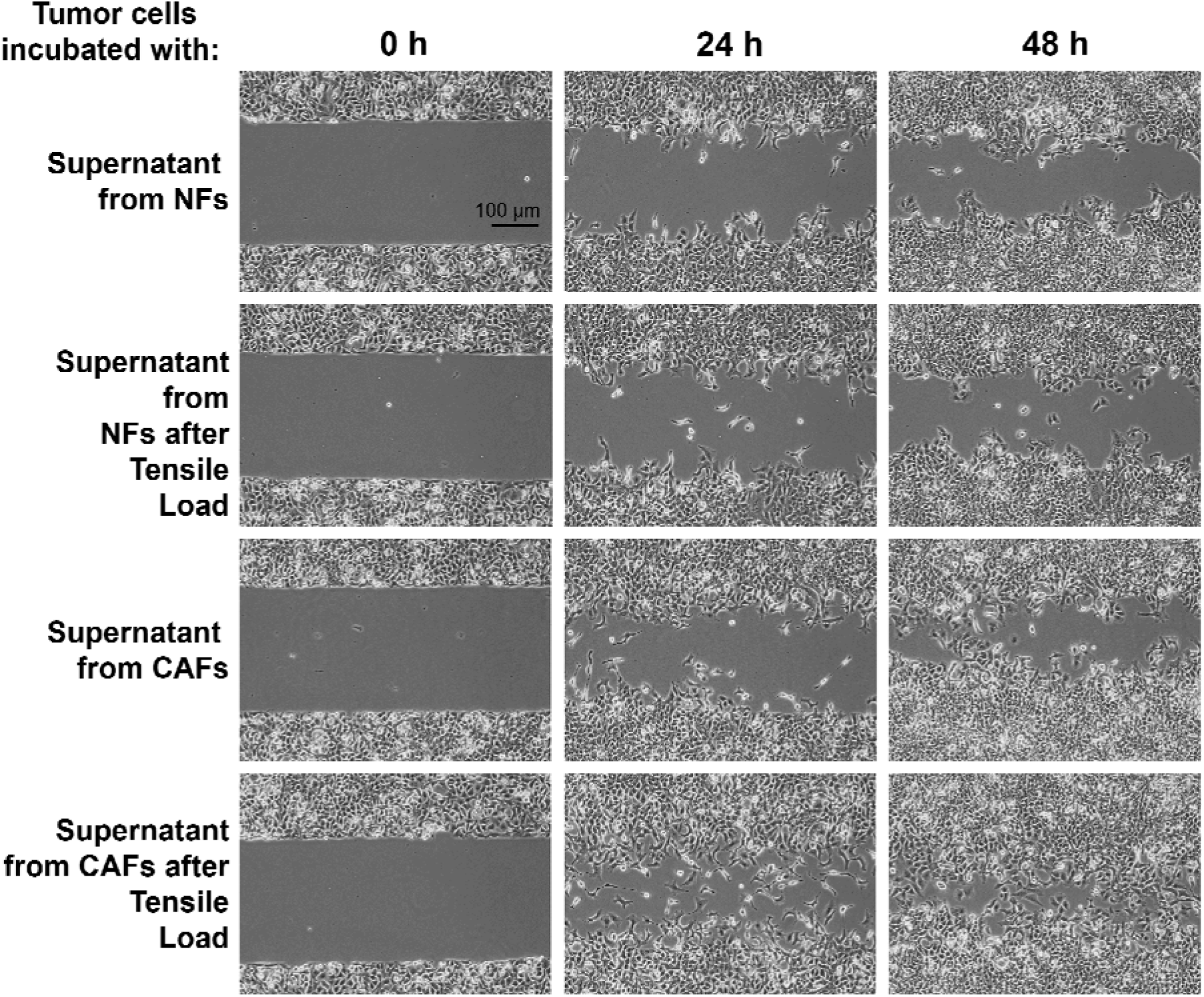
Representative Depiction of Tumor Cell Gap Closure after Incubation with Supernatant from NFs and CAFs of one Set of Patient Samples following Exposure to Tensile Load.

## Discussion

Currently, limited data exists on the mechanical properties and responses of CAFs under tensile forces, making this study a significant contribution to understanding the biomechanical aspects of oral tumor progression [39]. To the best of our knowledge, this is the first study to investigate the effects of tensile mechanical forces on oral CAFs and NFs, highlighting their distinct behaviour and roles within the TME. Our findings demonstrate that tensile strain modulates CAF behaviour, including gap closure, and potential paracrine effects, thereby providing novel insights into biomechanical interactions relevant to OSCC progression.

Mechanical and oxidative stress, together with matrix stiffness, are well-established modulators of cancer progression and complicate the prediction of clinical outcomes [40, 41]. Internal and external physical forces can influence invasion and metastasis by altering ECM stiffness, inducing jamming or confinement phenomena, and affecting interstitial fluid pressure [42]. In the present study, we focused on cyclic tensile strain as a reductionist and highly controlled *in vitro* stimulus. This choice was primarily driven by feasibility and standardisation using the Flexcell® system, enabling reproducible loading across multiple patient-matched primary CAF/NF pairs. At the same time, tensile deformation is biologically plausible in the oral cavity during repetitive stretching associated with swallowing, chewing, and speech, and may also contribute to stromal deformation at the invasive front as tumour growth and matrix remodelling impose tension within the surrounding extracellular matrix. The study of mechanical forces, particularly in the context of tumor–stroma interactions, is gaining increasing significance and is not restricted to the oral cavity [43, 44]. Importantly, tensile strain captures only one component of the *in vivo* mechanical milieu; compression and shear may elicit distinct cellular programmes and should be addressed in future studies. Interactions between tumor cells and stromal components, particularly CAFs, contribute to tumor heterogeneity, drug resistance, and metastatic capacity [45, 46]. ECM stiffening mediated by CAFs - largely through LOX-dependent collagen cross-linking - plays a central role in promoting invasiveness [47]. Mechanotransduction pathways such as YAP/TAZ, integrins, and cGAS-STING link these mechanical cues to alterations in stromal and cancer cell phenotypes [11, 20]. Furthermore, CAFs have been implicated in immune evasion, ECM remodeling, and resistance to therapy, underscoring their importance in tumor progression [48]. There is also evidence that CAFs facilitate cancer cell escape via mechanical interactions within the ECM [46].

Our study corroborates these observations by showing that CAFs exhibit superior migratory capabilities relative to NFs, with the most pronounced effects observed under tensile stimulation. Although little is known about fibroblast responses to tensile forces at the cellular level, existing evidence indicates that fibroblasts act as highly sensitive modulators of ECM mechanics. Bertillot et al. demonstrated that compressive stress generated by tumor expansion may trigger fibroblasts to form a capsule around the tumor [49]. Other studies have shown that mechanical stress at the invasion front can induce fibroblast-to-myofibroblast transition independently of exogenous TGF-β, accompanied by increased ECM deposition [50, 51]. These findings align with reports that CAFs promote directed migration of cancer cells through ECM remodelling and immune modulation [41], with mechanical forces acting as potent regulators of cellular behavior [39]. ECM synthesis, stiffening, reorganization, and guidance of cancer cell migration are proposed as major CAF-driven processes [52], and CAF-mediated immune suppression further contributes to tumor progression [53–55]. Nonetheless, the precise mechanisms underlying CAF mechanoactivation remain incompletely understood [56].

In our study, CAF proliferation under tensile stress showed minor changes that did not reach statistical significance, suggesting that their primary mechanoresponsive adaptation favours gap closure and ECM modulation rather than cell division. Mechanical loading also appeared to induce a metabolic shift in CAFs, reflected by a slight reduction in metabolic activity in stimulated CAFs. This may represent a redistribution of metabolic resources toward cytoskeletal reorganization, ECM remodelling, or paracrine signalling, processes known to be energetically demanding and central to CAF function. The multifaceted role of CAFs in shaping the TME, including angiogenesis and modulation of immune cell phenotypes [27, 57–60], supports this interpretation and emphasizes their adaptability to mechanical cues. The paracrine effects of CAFs on tumor cell gap closure became progressively more evident with increasing incubation time. Although only a non-significant trend was observed at 24 h, significant differences emerged at later time points (48 h, 72 h, 96 h), especially when A549 cells were exposed to supernatants from mechanically stimulated CAFs. This time-dependent increase suggests either cumulative secretion of pro-migratory factors or a delayed tumor cell response. In contrast, supernatants from mechanically stimulated NFs showed a reduced ability to support tumor cell gap closure, indicating distinct mechanosensitive phenotypes of CAFs versus NFs. These findings align with previous studies showing that CAF-conditioned media can promote migration and EMT via factors such as POSTN, SDF-1, PAI-1, TGF-β1, and via activation of chemokine receptors CCR2, CCR5, CXCR1/2, or Ras-associated pathways [61, 62]. Identifying the specific mediators responsible for the effects observed here warrants future investigation. Potential candidates include cytokines (e.g., HIF-1α), chemokines (e.g., CXCL12), and angiogenic factors (e.g., VEGFA, bFGF) [63–65]. The data presented highlight the amplification of CAF paracrine activity under tensile loading and underscore the relevance of biomechanical forces in shaping secretory phenotypes. However, we note that the conditioned-media experiments were performed using A549 cells as a standardized gap-closure readout model and not an OSCC cell line. Therefore, the observed effects should be interpreted as a general pro-migratory paracrine influence of oral CAF supernatants under the conditions tested, while OSCC-specific tumour-stroma interactions require confirmation in oral cancer cell lines and more complex co-culture/3D models.

Mechanotransduction in fibroblasts has been associated with several signalling axes, including YAP/TAZ-related transcriptional activity [45, 66], integrin-FAK signalling, and innate immune sensing pathways such as cGAS-STING [67]. These cascades have been implicated in ECM remodelling, immune evasion, and enhanced cellular migration and are known to respond dynamically to tensile forces and matrix stiffness [68, 69]. Previous multi-omic studies across several tumor entities have shown that these pathways are consistently upregulated in CAFs [70, 71]. However, these pathways were not directly assessed in the present study. Therefore, we interpret our findings as functional observations (gap closure and conditioned-media effects) and consider pathway involvement as a hypothesis that requires validation by dedicated signalling readouts (e.g., phospho-protein and intracellular localization analyses, knock-in and knock-out studies, transcriptomic/proteomic profiling).

Mechanical stimuli within the TME are increasingly recognized as key regulators of cell behaviour and tumor progression [72]. Our findings demonstrate that cyclic tensile strain is associated with increased gap closure and enhanced pro-migratory paracrine effects of CAFs under the conditions tested. While CAF-mediated stromal remodelling has been linked to epithelial plasticity in other settings [73, 74], EMT-related endpoints were not assessed here. Future studies should therefore evaluate whether mechanically stimulated CAF phenotypes translate into epithelial changes in OSCC-specific co-culture or 3D models.

Clinically, these findings may be particularly relevant for OSCC arising in anatomical sites subject to frequent mechanical deformation (e.g., the tongue, cheek, and floor of the mouth). Our data support the concept that tensile cues can modulate CAF functional behaviour and secretory activity *in vitro*. Whether such responses can be exploited therapeutically remains speculative and will require mechanistic validation and *in vivo* confirmation.

This study has several limitations. First, the cohort size was limited to five patient-matched CAF/NF pairs. Given the inherent heterogeneity of primary human fibroblast cultures, statistical power is limited, and comparisons with high variance or borderline p-values should be interpreted cautiously; therefore, the present dataset is best considered descriptive and hypothesis-generating. Second, proliferation was not pharmacologically inhibited during the scratch assays, and it was quantified only up to 48 h; therefore, the wound-healing readout should be interpreted conservatively as gap closure (combined migration and proliferation), at later time points (72-96 h). The absence of significant proliferation differences up to 48 h argues against a major early proliferation-driven explanation of the observed group differences [75], but it does not exclude a proliferative contribution at the later time points. Third, CAF/NF identity was established primarily by a patient-matched multi-marker qPCR panel, whereas protein-level α-SMA confirmation in vitro was performed in one representative CAF/NF pair only. Given the recognized of CAF populations [75] and the existence of α-SMA-low oral CAF states [76], this protein-level validation should be viewed as supportive rather than exhaustive; broader protein-level phenotyping across all isolates would strengthen future studies. Also, no pathway-level experiments were performed; therefore, mechanistic conclusions regarding YAP/TAZ-, integrin-FAK-, or cGAS-STING-related signalling cannot be drawn from the present dataset. In addition, the use of A549 cells for conditioned-media experiments represents a non-OSCC-specific tumour readout and limits the direct generalizability of these paracrine findings to oral oncology. Furthermore, the applied uniaxial cyclic tensile strain represents only one component of oral biomechanics and does not model relevant compressive or shear forces (e.g., during mastication, tongue pressure, denture-related loading, or tumor mass expansion). In addition, the selected strain amplitude and duration were based on literature precedent and feasibility rather than direct *in vivo* measurements of strain magnitudes in OSCC tissues.

As an *in vitro* analysis, the present study cannot fully recapitulate the complexity of the *in vivo* TME. More complex platforms, such as 3D cultures, organotypic models, or tumour-stroma co-culture systems, will be required to assess epithelial responses and to integrate additional force modalities in a more physiologically relevant context. Finally, although EMT is discussed as a potential downstream phenomenon in OSCC biology, EMT-related endpoints were not assessed in this study and cannot be inferred from the present dataset. Comparative analyses of mechanical properties and traction forces among CAFs, cancer cells, and precursor phenotypes such as NFs or epithelial cells could provide further insight into mechanoadaptive hierarchies. In addition, simulation-based modelling approaches may help reproduce clinical observations and guide hypothesis generation. Complementary biophysical measurement modalities, including surface pressure assessment or electrical conductivity measurements, could detect stiffness and ECM alterations and may have translational potential in diagnostic settings.

### Conclusions

This pilot study provides first evidence that cyclic tensile strain is associated with increased gap closure and enhanced pro-migratory paracrine effects of primary oral CAFs compared with patient-matched normal fibroblasts on tumor cells. Incorporating defined mechanical cues into *in vitro* models may improve the physiological relevance of stromal research in OSCC. Larger cohorts and pathway-level analyses in advanced co-culture or 3D systems will be required to define the molecular mechanisms and translational significance of these observations.

## List of Abbreviations

α-SMA: alpha-smooth muscle actin
BCS: bovine calf serum
bFGF: basic fibroblast growth factor
BSA: bovine serum albumin
CAF: cancer-associated fibroblast
CD31: cluster of differentiation 31
CD45: cluster of differentiation 45
COL1A2: collagen type I
cGAS: cyclic-GMP-AMP synthase
CXCL-12: C-X-C motif chemokine ligand 12
DAB: 3,3 ′ -Diaminobenzidine
ECM: extracellular matrix
EMT: epithelial–mesenchymal transition
EPCAM: epithelial cell adhesion molecule
FAP: fibroblast activation protein
FCS: fetal calf serum
FSP-1: fibroblast-specific protein-1 = S100A4
G2: moderately differentiated tumor
HBA1: hemoglobin alpha chain CD31 gene
HIF-1α: hypoxia-inducible factor 1-alpha
HNSCC: head and neck squamous cell carcinoma
HRP: horseradish peroxidase
IFP: interstitial fluid pressure
L0: no lymphatic invasion
LOX: lysyl oxigenase
NF: normal fibroblast
NSCLC: non-small-cell lung cancer
OSCC: oral squamous cell carcinoma
P1-5: Patient 1-5
PBS: phosphate-buffered saline
PDGF: platelet-derived growth factor
PDGFRA: platelet-derived growth factor receptor A CD140A
PDGFRB: platelet-derived growth factor receptor B CD140B
PDPN: podoplanin
pN: pathological lymph node status
Pn0: no perineural invasion
pT: pathological tumor size
PTPRC: protein tyrosine phosphatase receptor type C
qPCR: real-time quantitative PCR
R0: resection with clear margins
S100A4: S100 calcium binding protein A4 FSP-1
SD: standard deviation
STING: stimulator of interferon gene
TBST_20_: Tris-buffered saline with 0.05% Tween-20
TFRC: transferrin receptor CD71
TME: tumor microenvironment
TNM: Tumor, Node, Metastasis
V0: no vascular invasion
VEGFA: vascular endothelial growth factor A
VIM: vimentin
YAP/TAZ: yes-associated protein / transcriptional coactivator with PDZ-binding motif

## Declarations

### Ethics approval and consent to participate

Fibroblasts from tumor tissue and healthy mucosa were isolated from patients who received treatment at the Department of Oral and Maxillofacial Surgery - Plastic Surgery of the Johannes Gutenberg-University Mainz. All patients provided signed informed consent before participating in the study. The State Medical Association of Rhineland-Palatinate Ethics Committee approved this study (ethics vote: 2022-16424_3; year of ethics vote: 2022).

### Consent for publication

Not applicable.

### Availability of data and materials

The datasets used and analysed during the current study are available from the corresponding author on reasonable request.

### Competing interests

The authors declare that they have no competing interests.

## Funding

The study was financed by intramural funding and a grant provided by the Foundation Tumor Research Head and Neck, Wiesbaden, Germany. The foundation is a non-profit organization. The funders played no role in the experiment design, execution, analysis or preparation of the publication.

### Authors’ contributions

Conceptualization: NWI, JD, JB, PWK; methodology: NWI, JB, JD, AVBN, SK, VL, SZ; experiments: NWI, SK, AVBN, VL, SZ; validation: NWI, JM, AVBN, JUM, JD, JB, PWK, VL, SZ; formal analysis: NWI, PWK; data curation: NWI, JUM; writing—original draft preparation: NWI, PWK; writing-review and editing: NWI, SK, AVBN, JUM, JD, JB, PWK, VL, SZ; visualization: NWI; supervision: JD, JB, PWK; project administration: NWI, PWK.

All authors have read and agreed to the final version of the manuscript.

## Acknowledgements

We are greatly indebted to Simone Mendler and Rita Gieringer for their excellent technical support. We also want to thank Jutta Goldschmitt, Jutta Bühler, and Christina Babel for their support in the Department of Oral and Maxillofacial Surgery’s laboratory. Furthermore, we thank Antonietta Valentino for her outstanding technical support in the Periodontology and Operative Dentistry laboratory, and Gargi Nayak for his help with the manuscript preparation. Furthermore, we thank the team of the tissue bank of the University Medical Center Mainz for providing and staining the tissue samples in accordance with the regulations of the tissue bank, especially Silke Mitschke and Bonny Adami.

## Notes

### Competing Interest Statement

The authors have declared no competing interest.

